# Genome-wide patterns of population structure and linkage disequilibrium in farmed Nile tilapia (*Oreochromis niloticus*)

**DOI:** 10.1101/519801

**Authors:** Grazyella M. Yoshida, Agustín Barria, Katharina Correa, Giovanna Cáceres, Ana Jedlicki, María I. Cadiz, Jean P. Lhorente, José M. Yáñez

**Affiliations:** Facultad de Ciencias Veterinarias y Pecuarias, Universidad de Chile, Santiago, Chile; Benchmark Genetics Chile, Puerto Montt, Chile; Nucleo Milenio INVASAL, Concepción, Chile

**Keywords:** effective population size, genomic prediction, GWAS, *Oreochromis niloticus*, population structure

## Abstract

Nile tilapia (*Oreochromis niloticus*) is one of the most produced farmed fish in the world and represents an important source of protein for human consumption. Farmed Nile tilapia populations are increasingly based on genetically improved stocks, which have been established from admixed populations. To date, there is scarce information about the population genomics of farmed Nile tilapia, assessed by dense single nucleotide polymorphism (SNP) panels. The patterns of linkage disequilibrium (LD) may affect the success of genome-wide association studies (GWAS) and genomic selection and can also provide key information about demographic history of farmed Nile tilapia populations. The objectives of this study were to provide further knowledge about the population structure and LD patterns, as well as, estimate the effective population size (*N*_*e*_) for three farmed Nile tilapia populations, one from Brazil (POP A) and two from Costa Rica (POP B and POP C). A total of 55, 56 and 57 individuals from POP A, POP B and POP C, respectively, were genotyped using a 50K SNP panel selected from a whole-genome sequencing (WGS) experiment. Two principal components explained about 20% of the total variation and clearly discriminated between the three populations. Population genetic structure analysis showed evidence of admixture, especially for POP C. The contemporary *N*_*e*_ values calculated based to LD values, ranged from 71 to 141. No differences were observed in the LD decay among populations, with a rapid decrease of *r*^*2*^ when increasing inter-marker distance. Average *r*^*2*^ between adjacent SNP pairs ranged from 0.03 to 0.18, 0.03 to 0.17 and 0.03 to 0.16 for POP A, POP B and POP C, respectively. Based on the number of independent chromosome segments in the Nile tilapia genome, at least 4.2 K SNP are required for the implementation of GWAS and genomic selection in farmed Nile tilapia populations.

## Introduction

Nile tilapia (*Oreochromis niloticus*) is one of most important farmed fish species worldwide. Breeding programs established since the 1990’s have played a key role in improving commercially important traits and expanding Nile tilapia farming. The Genetically Improved Farmed Tilapia (GIFT) is the most widespread tilapia breeding strain (Lim and Webster, 2006), which has been introduced to several countries in Asia, Africa and Latin America (Gupta and Acosta, 2004). The genetic base of GIFT was established from eight African and Asian populations, and after six generations of selection, the genetic gains ranged from 10 to 15% per generation for growth-related traits (Ponzoni et al., 2011), providing evidence that selective breeding using phenotype and pedigree information can achieve high and constant genetic gains (Gjedrem and Rye, 2018).

The recent development of single nucleotide polymorphism (SNP) panels for tilapia (Joshi et al., 2018; Yáñez et al., submitted) will provide new opportunities for uncovering the genetic basis of relevant traits through GWAS and improving traits that are difficult or expensive to measure in selection candidates (e.g. fillet yield and diseases resistance) by the use of genomic selection (Meuwissen et al., 2001). As has been demonstrated for different traits in salmonid species, the incorporation of genomic prediction into breeding programs is expected to increase the accuracy of breeding value predictions, compared to pedigree-based methods (Bangera et al., 2017; Barria et al., 2018b; Correa et al., 2017; Sae-Lim et al., 2017; Tsai et al., 2016; Vallejo et al., 2018; Yoshida et al., 2017, 2018).

Genomic studies exploit the linkage disequilibrium (LD) between SNP and quantitative trait locus (QTL) or causative mutation. Thus knowing the extent and decay of LD within a population is important to determine the number of markers that are required for successful association mapping and genomic selection (Brito et al., 2015; de Roos et al., 2008; Khatkar et al., 2008; Porto-Neto et al., 2014). To achieve a low LD level requires a higher marker density to enable markers to capture most of the genetic variation in a population (Khatkar et al., 2008). The demographic history of farmed fish populations is one of factors that can affect the extent and decay of LD, which in turn may affect the success of GWAS and genomic prediction. In addition, LD patterns provide relevant information about past demographic events including response to both natural and artificial selection (Slatkin, 2008). The LD throughout the genome, besides reflecting the population history, provides insight about the breeding system and pattern of geographic subdivision, which can be explored to study the degree of diversity in different populations.

To date, the most widely used measures of LD between two loci are r^2^ and Lewontin’s D’ (commonly named D’). Values lower than 1 for D’ indicate loci separation due to recombination, while D’ = 1 indicates complete LD between loci, i.e. no recombination. However, this parameter is highly influenced by allele frequency and sample size. Thus, high D’ estimations are possible even when loci are in linkage equilibrium (Ardlie et al., 2002). Therefore, LD measured as the squared correlation (r^2^) between two loci is suggested as the most suitable measurement for SNP data (Pritchard and Przeworski, 2001).

LD patterns have been widely studied in different livestock species, such as sheep (Prieur et al., 2017), goats (Mdladla et al., 2016), pigs (Ai et al., 2013), beef (Espigolan et al., 2013; Porto-Neto et al., 2014) and dairy cattle (Bohmanova et al., 2010). In aquaculture, recent studies have aimed at characterizing the extent and decay of LD in farmed species, such as Pacific white shrimp (Jones et al., 2017), Pacific oyster (Zhong et al., 2017), rainbow trout (Rexroad and Vallejo, 2009; Vallejo et al., 2018), coho salmon (Barria et al., 2018a) and Atlantic salmon (Barria et al., 2018c; Gutierrez et al., 2015; Hayes et al., 2006; Kijas et al., 2016). However, to date there is scarce information about population genomic structure and LD in farmed Nile tilapia assessed by the use of dense SNP panels. For instance, the assessment of LD patterns in Nile tilapia is still limited to a few studies in which either a small number of markers (14 microsatellites) (Sukmanomon et al., 2012) and individuals (4 to 23 samples) (Hong Xia et al., 2015) have been used. The objectives of the present study were to (i) estimate the population structure and genetic differentiation; (ii) assess the genome-wide levels of LD and (iii) determine the effective population size among three Nile tilapia breeding populations established in Latin America.

## Methods

### Populations

Samples were obtained from three different commercial breeding populations established in Latin America, originating from admixed stocks imported from Asia and genetically improved for growth rate for more than 20 generations. Population A (POP A) was obtained from the AquaAmerica (Brazil) breeding population. This population was imported from Malaysia in 2005 for breeding and farming purposes, with a genetic origin from the GIFT strain. POP B and POP C populations were obtained from Aquacorporación Internacional (Costa Rica). The POP B breeding population is a mixture of the GIFT strain, POP C and strains from Egypt and Kenya. The POP C breeding population represents a combination of genetic material from Israel, Singapore, Taiwan and Thailand from the GIFT strain in the Philippines. Therefore, the three breeding populations are considered recently admixed populations; which are directly or indirectly related to the GIFT strain and have been artificially selected to improve growth-related traits. The average relatedness between individuals, within each population, was estimated using Plink v1.90 (Purcell et al., 2007).

### Genotyping

The genotypes were selected from a whole-genome sequencing experiment aimed at designing a 50K SNP Illumina BeadChip, which is described in detail in Yáñez et al. (submitted). Briefly, a total of 59, 126 and 141 individuals were fin-clip sampled for POP A, POP B and POP C, respectively. Genomic DNA was purified from all the samples using the DNeasy Blood & Tissue Kit (QIAGEN) according to the manufactured’s protocol. Whole-genome sequencing was performed using multiplexing of four bar-coded samples per lane of 100bp paired-end in the Illumina HiSeq 2500 machine. The sequences were trimmed and aligned against the genome assembly O_niloticus_UMD1 (Conte et al., 2017). About 36 million polymorphic sites were discovered after variant calling using the Genome Analysis Toolkit GATK (McKenna et al., 2010). A list of 50K SNP were selected based on quality of genotype and site, number of missing values, minor allele frequency (MAF), unique position in the genome and even distribution across the genome as described by Yáñez et al (submitted). Furthermore, genotype quality control (QC) was performed within each population excluding SNPs with MAF lower than 5%, Hardy-Weinberg Equilibrium P-value < 1e-06, and missing genotype higher than 70%. Animals with a genotype call rate below 95% were discarded. Afterwards, to use a similar sample size among populations, animals from POP B and POP C with high identical by descent (IBD) were excluded. The SNP markers used in the subsequent analyses are those common among the three populations after QC.

### Population structure

We used the software Plink v1.09 (Purcell et al., 2007) to calculate the heterozygosity observed (H_o_) and expected (H_e_) for the three populations and for genetic differentiation through principal component analysis (PCA). The results of the first two PCAs were plotted along two axes using R scripts (R Core Team, 2016). Additionally, the population structure was examined using a hierarchical Bayesian model implemented in STRUCTURE software v.2.3.4 (Pritchard et al., 2000). We used three replicates of K value ranging from 1 to 10, a burn-in of 20,000 iterations and running of 50,000. To choose the best K value we computed the posterior probability of each K as suggested by Pritchard et al., (2000).

### Estimation of linkage disequilibrium and effective population size

We used the Pearson’s squared correlation coefficient (r^2^) to estimate the LD between each pair of markers separated by an inter-marker distance between 0 and 10 Mb for each population. We used Plink v1.09 (Purcell et al., 2007) using the parameters --ld-window-kb 10000 and --ld-window-r2 set to zero to calculate the LD between all pairs of SNPs on each chromosome. The extent and decay of the LD, for each population, were visualized by plotting moving average LD window of 10 Mb along inter-marker distances.

We used the software SNeP v1.1 (Barbato et al., 2015) to estimate the historical effective population size (*N*_e_). Considering the LD within each population, *N*_*e*_ was estimated using the following equation proposed by Corbin et al., (2012):

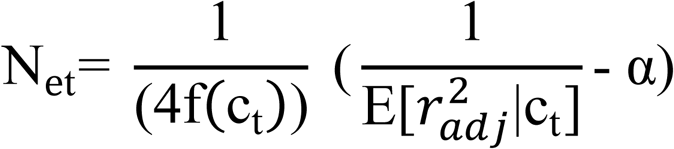

where *N*_*et*_ is the effective population size *t* generations ago, *c*_*t*_ represents the recombination rate *t* generations ago, which is proportional to the physical distance between SNP pair markers, *r*^*2*^_*adj*_ is the estimated LD corrected for sample size and α is the adjustment for mutation rate (α = 2, indicate the presence of mutation). We grouped the data in 30 distance bins of 50 Kb each. Based on the relatively small number of SNP per chromosome, *N*_e_ per chromosome was calculated using harmonic mean (Alvarenga et al., 2018). Using the LD method, we calculated the contemporary population size using the software NeEstimator v2.01 (Do et al., 2014), using a non-random mating model and a critical value (Pcrit) of 0.05.

## Results

### Quality control

Out of the initial 50K SNP, a total of 31,176 markers were shared among the three populations after QC criteria. The MAF < 0.05 excluded the higher number of SNPs (~ 9.7K on average). After QC, all three populations showed a similar mean MAF value of 0.26 ± 0.13. The proportion of SNPs for each MAF class was very similar among the populations (**Figure 1**). The lower (~ 0.15) and higher (~ 0.25) proportion of SNP were observed in the MAF classes ranging from 0.05 to 0.09 and 0.10 to 0.19, respectively. To conduct downstream analysis, we used 55, 56 and 57 animals for POP A, POP B and POP C, respectively. These individuals represent the animals with the lowest levels of IBD, among all sequenced animals, within each population. Average relatedness among the selected individuals was equal to zero within populations.

**Figure 1.**
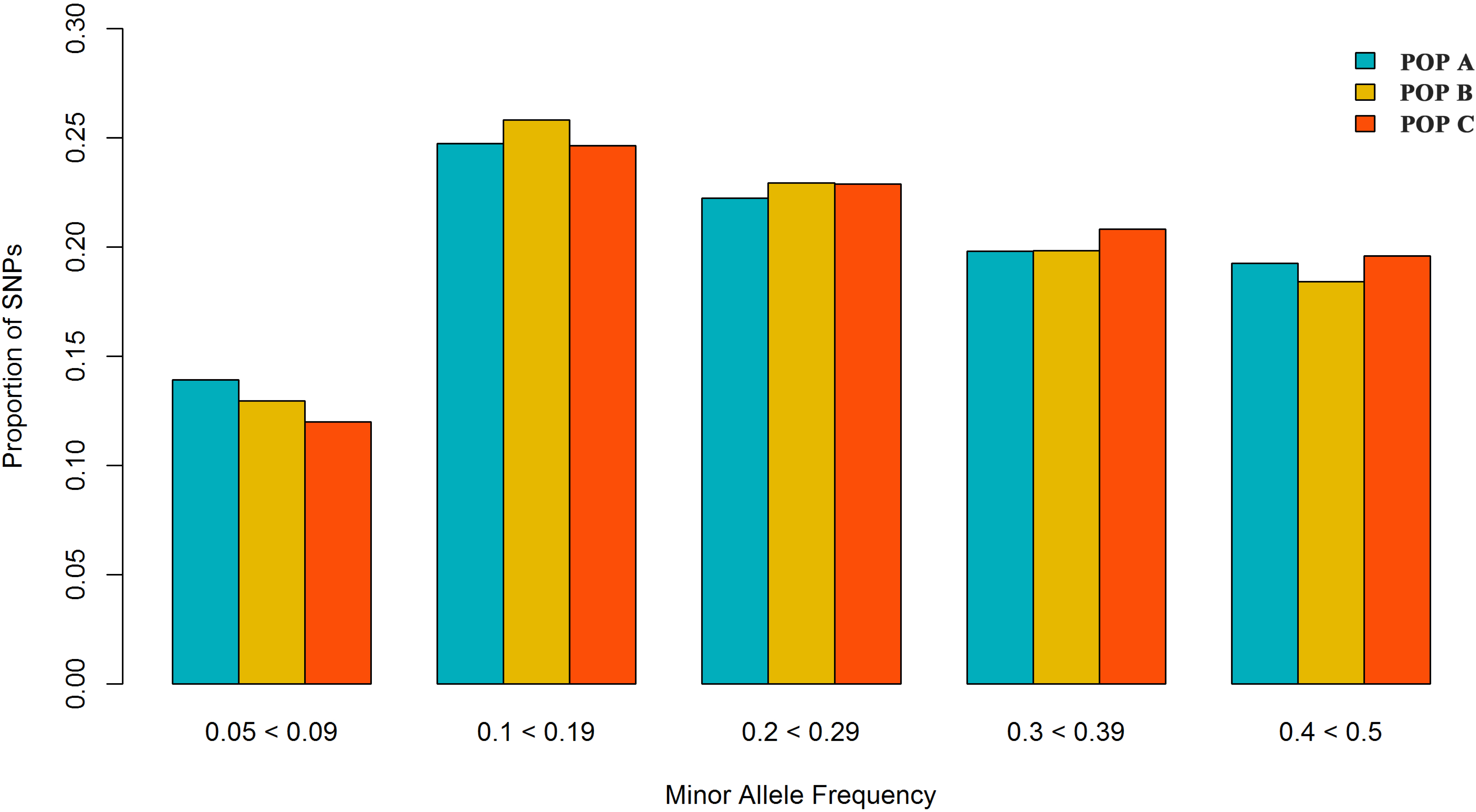
Proportion of SNPs for different minor allele frequency for three Nile tilapia populations.

### Population structure

Upon plotting the first two eigenvectors on the PCA plot, the three populations were stratified based on the single dimensional variation between them. The first two principal components together accounted for 20.0% of the genetic variation, clearly revealing the three different populations (**Figure 2**). PCA1 discriminates between populations from Brazil and Costa Rica and accounted for 11.3% of the total genetic variation. The second principal component explains 8.7% of the total variance and separated the two populations from Costa Rica into two different clusters. To assess the genetic diversity among populations, we calculated the observed/expected heterozygosity ratio (H_o_/H_e_). We found values of 0.23/0.37, 0.26/0.35 and 0.24/0.35 for POP A, POP B and POP C respectively.

**Figure 2.**
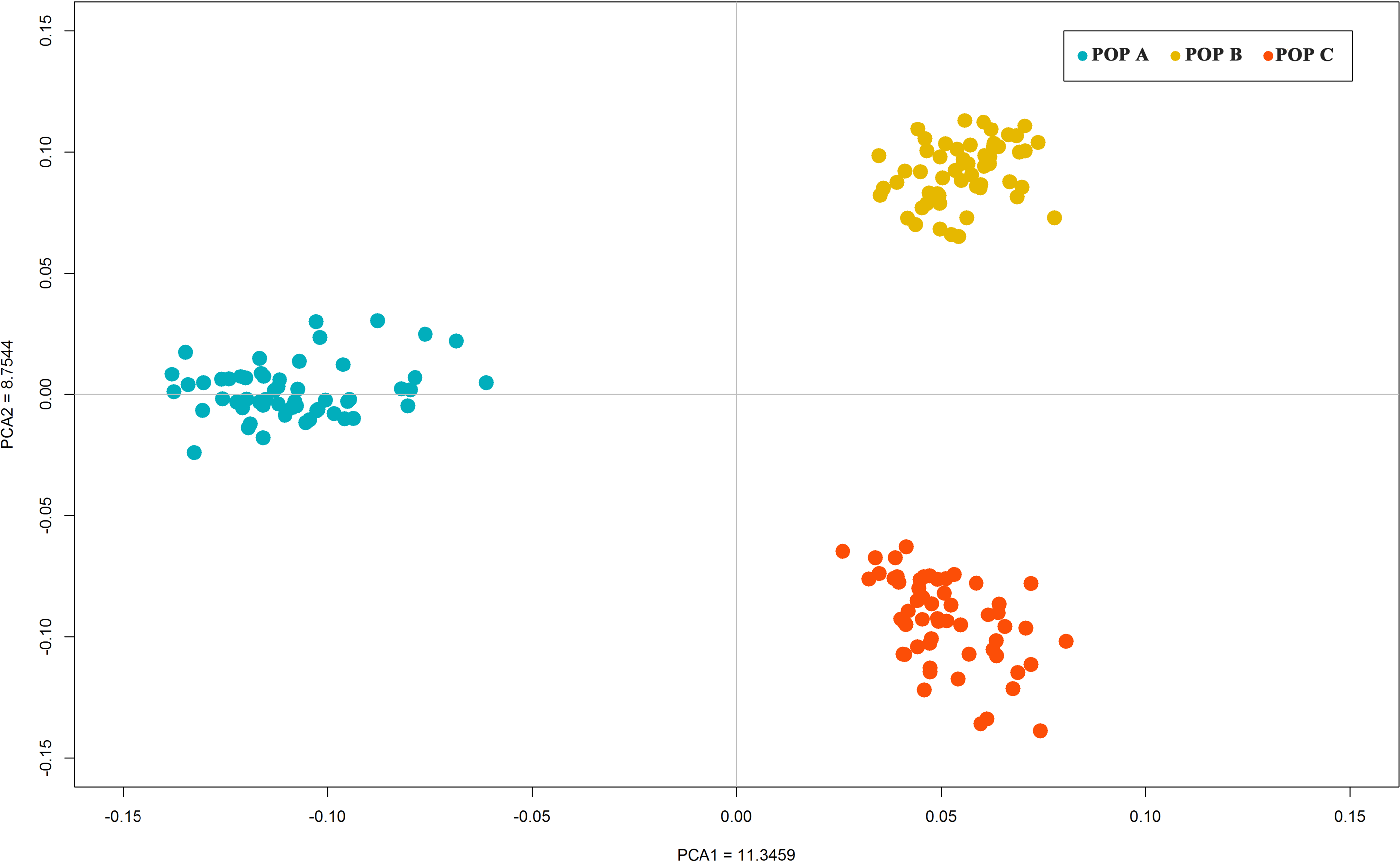
Principal component analysis revealing genetic differentiation of three Nile tilapia populations using autosomal genotypic data.

In the admixture analysis, by computing the posterior probabilities of each K, the best result was K = 9. Different genomic clustering levels and a high level of admixture was observed for the three populations studied. POP B and POP C showed higher similarity between each other, than with POP A. For all populations some individuals contained a level of genetic variation from another population; however, POP B shared a high proportion (>0.25) of one subpopulation common to POP A (red color) and POP B (orange color) (**Figure 3**). STRUCTURE results evaluating K values from 2 to 10 are presented in **Supplementary File 1**.

**Figure 3.**
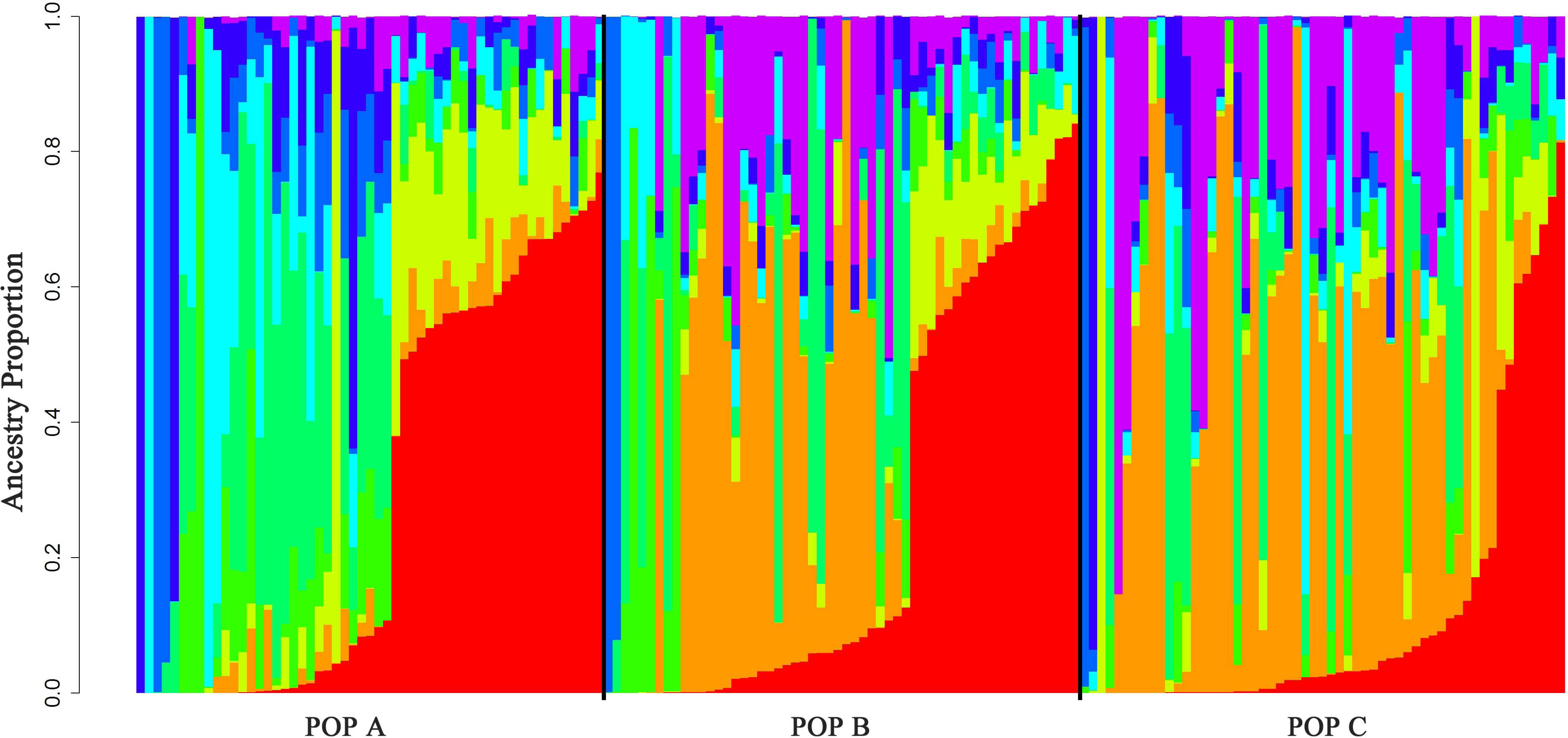
Admixture clustering of the three Nile population for K = 9. The animals are grouped by population and each individual is represented by a vertical bar. The gradient black lines delineate different populations under study.

### Estimation of linkage disequilibrium and effective population size

The overall mean LD between marker pairs measured by *r*^*2*^ was 0.06 for both POP A and POP C, and slightly lower (0.05) for POP B (**Table 1**). In general, the average LD among chromosomes ranged from 0.04 to 0.08 for all populations (**Table 1**). From one to 10,000 Kb, the average of *r*^*2*^ decreased with increasing physical distance between markers, from 0.18 to 0.03, 0.17 to 0.03 and 0.16 to 0.03 for POP A, POP B and POP C, respectively. In addition, the average LD decayed to less than 0.05 within 4 Mb (**Figure 4**), and this rate of decrease was very similar across all of the chromosomes for the three populations (**Supplementary File 2 to 4**).

**Table 1.** Number of SNPs, chromosome linkage group (LG) size, average linkage disequilibrium (r^2^) and effective population size (*N*_*e*_) for three Nile tilapia farmed populations.

| Chromossome |  |  | POP A |  | POP B |  | POP C |  |
| --- | --- | --- | --- | --- | --- | --- | --- | --- |
| ONil | Number of SNPs | Size (Mb) | $r^2$ mean ± SD | $N_e$ | $r^2$ mean ± SD | $N_e$ | $r^2$ mean ± SD | $N_e$ |
| 01 | 1,386 | 38.26 | 0.05±0.08 | 173 | 0.06±0.09 | 154 | 0.05±0.08 | 174 |
| 02 | 1,186 | 35.09 | 0.07±0.11 | 139 | 0.05±0.08 | 184 | 0.06±0.09 | 159 |
| 03 | 1,617 | 54.17 | 0.04±0.06 | 229 | 0.04±0.06 | 239 | 0.04±0.06 | 214 |
| 04 | 1,313 | 37.93 | 0.05±0.08 | 192 | 0.05±0.07 | 199 | 0.05±0.07 | 186 |
| 05 | 1,219 | 34.51 | 0.05±0.09 | 183 | 0.05±0.09 | 178 | 0.05±0.08 | 179 |
| 06 | 1,522 | 44.55 | 0.06±0.10 | 151 | 0.05±0.08 | 200 | 0.06±0.08 | 156 |
| 07 | 2,558 | 61.88 | 0.06±0.10 | 138 | 0.06±0.10 | 154 | 0.06±0.09 | 151 |
| 08 | 1,273 | 30.74 | 0.06±0.10 | 152 | 0.05±0.09 | 186 | 0.05±0.09 | 185 |
| 09 | 1,036 | 27.47 | 0.05±0.08 | 215 | 0.05±0.08 | 223 | 0.05±0.08 | 200 |
| 10 | 1,272 | 32.35 | 0.05±0.09 | 181 | 0.06±0.09 | 166 | 0.06±0.09 | 161 |
| 11 | 1,330 | 36.27 | 0.05±0.09 | 180 | 0.05±0.08 | 201 | 0.06±0.09 | 156 |
| 12 | 1,492 | 41.13 | 0.05±0.08 | 191 | 0.05±0.09 | 176 | 0.05±0.08 | 168 |
| 13 | 1,137 | 32.25 | 0.06±0.12 | 142 | 0.05±0.09 | 187 | 0.06±0.09 | 162 |
| 14 | 1,544 | 39.18 | 0.05±0.08 | 190 | 0.05±0.09 | 171 | 0.06±0.09 | 142 |
| 15 | 1,213 | 36.09 | 0.06±0.09 | 156 | 0.06±0.09 | 159 | 0.06±0.09 | 145 |
| 16 | 1,715 | 43.61 | 0.06±0.10 | 158 | 0.07±0.11 | 138 | 0.05±0.09 | 176 |
| 17 | 1,409 | 40.58 | 0.06±0.09 | 153 | 0.06±0.09 | 152 | 0.05±0.08 | 168 |
| 18 | 1,363 | 36.96 | 0.06±0.10 | 165 | 0.05±0.09 | 193 | 0.05±0.08 | 183 |
| 19 | 1,202 | 31.16 | 0.08±0.14 | 130 | 0.08±0.14 | 123 | 0.08±0.13 | 119 |
| 20 | 1,457 | 36.56 | 0.06±0.09 | 167 | 0.05±0.08 | 204 | 0.05±0.08 | 176 |
| 22 | 1,518 | 36.92 | 0.05±0.09 | 164 | 0.06±0.09 | 156 | 0.06±0.10 | 145 |
| 23 | 1,414 | 43.89 | 0.05±0.09 | 176 | 0.05±0.08 | 186 | 0.06±0.10 | 143 |
| **Mean** | **1,417** | **46.86** | **0.06±0.10** | **169** | **0.05±0.09** | **179** | **0.06±0.09** | **166** |

**Figure 4.**
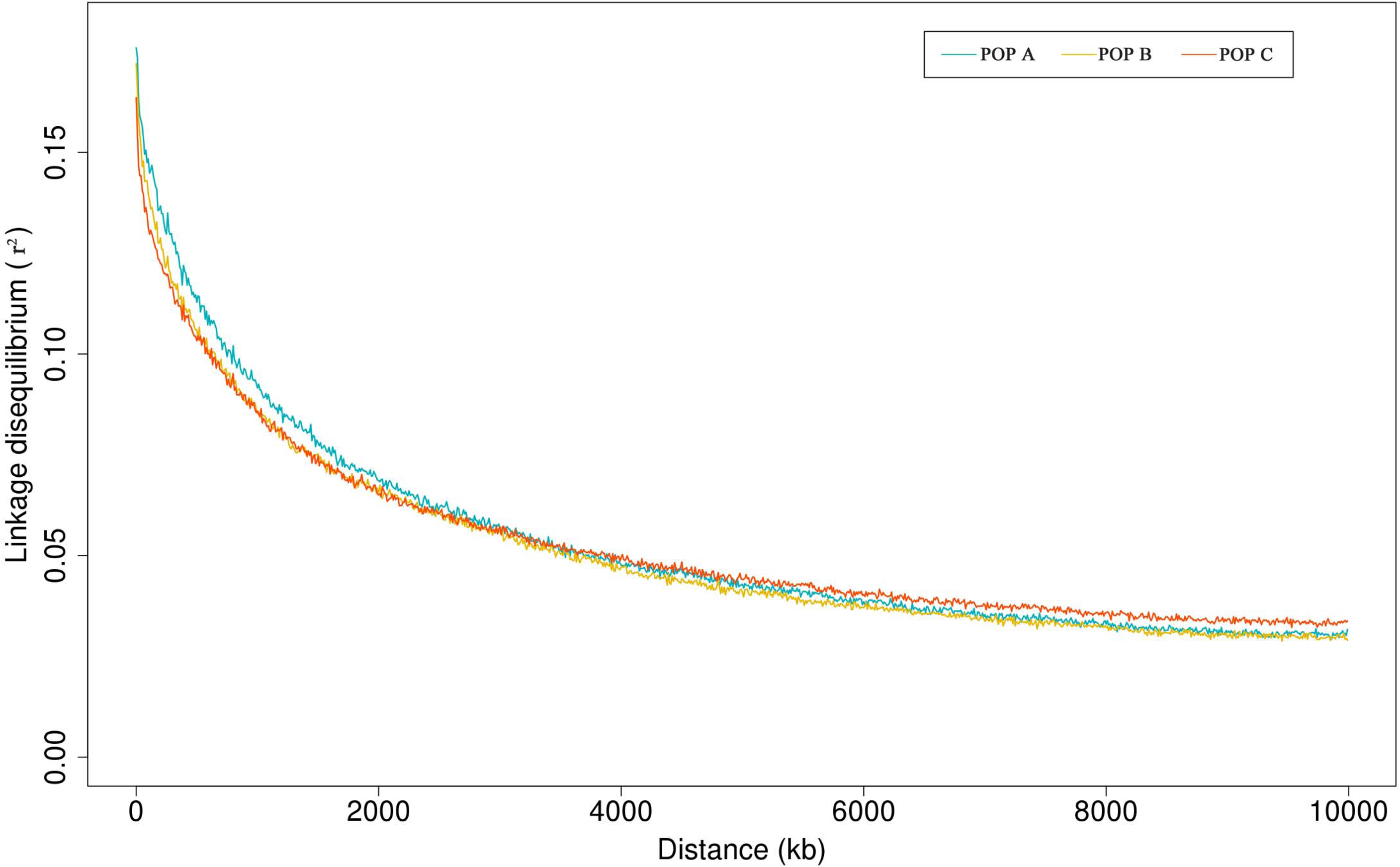
Average linkage disequilibrium (r^2^) decay by physical distance for three Nile tilapia populations.

**Figure 5** shows the historical *N*_e_ up to 1500 (A) and up to 120 (B) generations ago, respectively. The *N*_*e*_ values were lower in the recent past than the distant past. These values calculated at five generations ago were 80, 83 and 73 for POP A, POP B and POP C, respectively. The harmonic means for *N*_*e*_ at five to 1,500 generations ago was 169, 179 and 166 for POP A, POP B and POP C, respectively. In addition, the *N*_*e*_ varied among chromosomes, ranging from 119 to 239 (**Table 1**). Contemporary *N*_*e*_ calculated based on LD values were 141, 114 and 71 for POP A, POP B and POP C, respectively.

**Figure 5.**
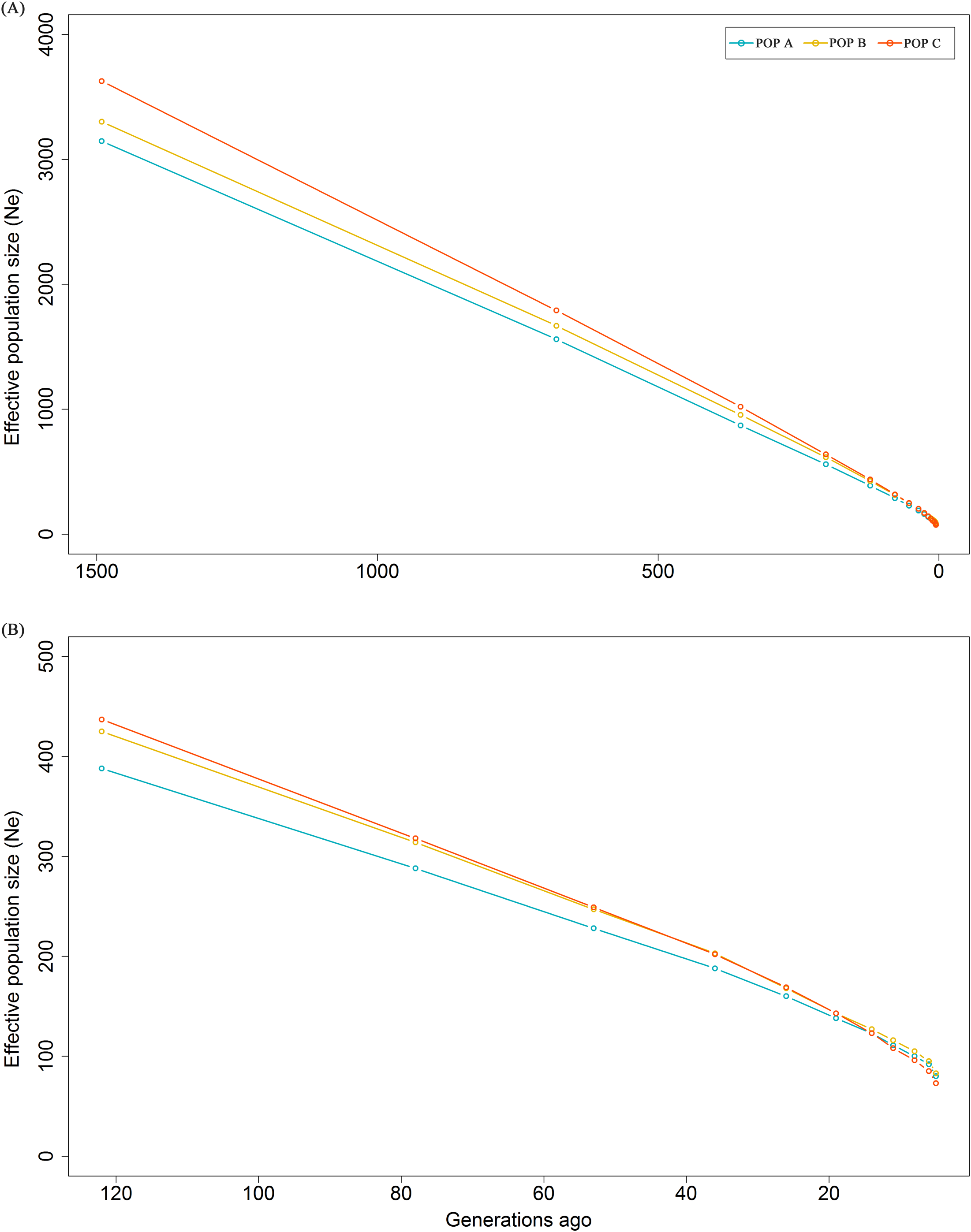
Effective population size (*N*_e_) from 1500 to 5 generations ago (a) and from 120 to 5 generations ago (b) based on linkage disequilibrium for three Nile tilapia populations.

## Discussion

### Genomic population structure

In the PCA, the first two principal components explained about 20% of the total genetic variation for the populations studied and clearly revealed three different clusters, corresponding to the three populations present in the dataset (**Figure 2**). In addition, the lowest values of H_o_ for POP A and POP C suggest a loss of genetic diversity due to found effect, effective population size, or the sample size used in this study.

The admixture results provided evidence of a recent mixture of different strains to conform highly admixture populations. Although the PCA demonstrates three distinct populations, the admixture analysis showed that, in fact, the three Nile tilapia populations studied are related through the common GIFT origin. The genetic differentiation among populations may have been be partly generated by genetic drift or founder effect events which can have a pronounced effect on allele frequencies (Allendorf and Phelps, 1980). Furthermore, the three populations have undergone artificial selection for the improvement of growth-related traits in different geographic locations exposing the populations to distinct environmental conditions and production systems (POP A and both POP B and C, in Brazil and Cost Rica, respectively). Previous studies suggest, the introgression of other tilapia species or strains, such as *O. mossambicus* or the Chitralada strain, into the GIFT stocks (McKinna et al., 2010; Sukmanomon et al., 2012; Xia et al., 2014).

### Linkage disequilibrium and effective population size

Evaluating the whole-genome LD within populations, may help to understand the different demographic processes experienced by these populations. These processes include admixture, mutation, founder effect, inbreeding and selection (Gaut and Long, 2003). This is the first study aimed at estimating the extent and decay of LD in farmed Nile tilapia populations established in Latin America (specifically, Brazil and Costa Rica), and artificially selected for growth-related traits. Previously, it has been suggested that due to high kinship relationships high levels of IBD among samples may inflate the LD estimate (Gutierrez et al., 2015). Thus, to reduce differences in average *r*^*2*^, we selected individuals based on low IBD values. Furthermore, to reduce bias sampling effect, we used a similar number of animals from each population. Another factor that may influence the LD estimate is MAF distributions (Espigolan et al., 2013). High frequency alleles result in less biased LD estimations. In the present study a small proportion of SNP (<15%) have MAF lower than 0.10 and low IBD values indicating an accurate estimation of LD. We used the squared correlation of allele frequencies as a measure of LD instead of |D’| to avoid overestimations of LD due the small sample size (Khatkar et al., 2008). The number of animals to estimate the LD accurately depends on the demographic and population history. POP A, POP B and POP C each had >55 individuals as suggested by Bohmanova et al., (2010) and Khatkar et al., (2008).

We observed on chromosome LG13 and LG19, a pool of *r*^*2*^ values >0.40 for pair-wise SNP at large distances (>7 Mb; Supplementary File 2-4), but a LD decline when physical distance between markers increases is expected. Incorrect position of SNPs on the reference genome or errors in the reference genome assembly may be resulted in errors in the estimates. Our study revealed that the LD level declined to 0.05 at 4000 Kb inter-marker distance and that decay patterns were similar between populations (**Figure 4**). A previous study conducted by Hong Xia et al., (2015) reported similar LD patterns for GIFT tilapia stocks collected from South Africa, Singapore and China. Using microsatellite loci Sukmanomon et al., (2012) estimated LD means in terms of disequilibrium coefficient (D’) of 0.05 for a GIFT population originally from the Philippines.

Although most of the time, differences between genomes, the quality control applied and population structure make LD comparison of different species inappropriate, here we used references from other farmed fish species because of the limited information that exists for this kind of study in tilapia. Thus, the Nile tilapia populations seems to present smaller levels of LD than other farmed fish populations (Barria et al., 2018b; Barria et al. 2018c; Gutierrez et al., 2015; Kijas et al., 2016; Vallejo et al., 2018). A likely explanation is due the diverse origin of the studied Nile tilapia populations, as was suggested for a Chilean farmed Atlantic salmon population with Norgewian origin (Barria et al. 2018c). In salmonids, some suggest admixture is a major factor contributing to long-range LD (Barria et al., 2018c; Ødegård et al., 2014; Vallejo et al., 2018). Our admixture results suggested, as expected, high evidences that the Nile tilapia populations have recent history of admixture with wild stocks or different strains, but not resulted in long-range LD.

For Nile tilapia, the effective population size could be the primary cause for the LD values in POP A, POP B and POP C. LD at a short distance is a function of effective population size many generations ago and LD at long distances reflect the recent population history. The LD estimation for POP A resulted in slightly different values at small distance compared to POP B and POP C, whereas at large distance the differences were more evident for POP C (**Figure 4**). These results were reflected in smaller *N*_*e*_ of many generations ago for POP A and smaller *N*_*e*_ in the recent past for POP C (**Figure 8**). However, the continuous reduction in the *N*_*e*_, regardless of population, was observed over the previous 1,500 generations (**Figure 5A**). The reduction of *N*_e_ can be considered an indicator of selection and has been suggested to be an important cause of LD (Pritchard and Przeworski, 2001) and the three populations in this study have been under genetic selection for some generations. Previously, similar values of *N*_*e*_ were estimated using pedigree information from a GIFT population from Malaysia (*N*_*e*_ = 88) and from Brazil (*N*_e_ = 95) (Ponzoni et al., 2010; Yoshida et al., submitted). Some suggest keeping *N*_*e*_ values ranging from 50 to 200 to ensure genetic variability in a long-term breeding population (Bijma, 2000; Smitherman and Tave, 1987). In contrast, a smaller *N*_*e*_ was found for for rainbow trout (Vallejo et al., 2018) and Atlantic salmon from North America, Europe (Barria et al., 2018c) and Tasmania (Kijas et al., 2016).

In summary, within tilapia populations, the LD values were very low even in short distances (*r*^*2*^ = 0.15 for markers spaced at 20-80 Kb). Similar values were found in humans (Ardlie et al., 2002; Reich et al., 2001), coho salmon (Barria et al., 2018a), some breeds of cattle (de Roos et al., 2008; Khatkar et al., 2008; Yurchenko et al., 2018), sheep (Alvarenga et al., 2018) and goats (Brito et al., 2015). Therefore, our LD results have several implications for future implementation of genomic tools in Nile tilapia. Both GWAS and genomic selection are dependent on LD extent to define the number of SNPs necessary to assure the causative mutation variance (Flint-Garcia et al., 2003) and to achieve a certain accuracy of genomic estimated breeding value (Meuwissen et al., 2001). Meuwissen (2009) suggested that to achieve accuracies of genomic breeding (GEBV) ranging from 0.88 to 0.93 using unrelated individuals; it is necessary to have *2N*_*e*_*L* number of individuals and *10N*_*e*_*L* number of markers, where L is the length of genome in Morgans. In our study, the contemporary *N*_e_ is 141, 114 and 71 for POP A, POP B and POP C, respectively, and the length of the genome is 14.8 Morgans (Conte et al., 2018). Thus the 11,000 to 21,000 markers are required for Nile tilapia populations. In contrast, Goddard (2009) suggested that accuracy of genomic prediction is highly dependent on the effective number of chromosome segments (*M*_*e*_*=4N*_*e*_*L)*. Having a number of independent, biallelic and additive QTL affecting the trait means we would need a smaller number of markers to achieve a high accuracy. Thus, the minimum number of markers for a high power genomic analysis should be at least, 8,300, 6,700 and 4,200 for POP A, POP B and POP C, respectively; numbers slightly lower than those suggested by Vallejo et al. (2018) and Barria et al. (2018a) for rainbow trout and coho salmon, respectively, using the same approach.

When the genome is sufficiently saturated with markers, the accuracy of GEBV may also depend on other factors such as the number of individuals genotyped and phenotyped in the training population and the heritability and number of loci affecting the trait (Daetwyler et al., 2008; Goddard, 2009). In preliminary studies of genomic prediction for Nile tilapia, we found high accuracies of GEBV (results not show) for complex traits, using a similar number of markers but a smaller number of animals suggested by Meuwissen (2009). However, this is our first genomic prediction analysis and we have still to test other experimental designs, marker density and methods to confirm the relationship between the number of markers and accuracy of GEBV. Once completed, it will be possible to cost-effectively include genomic information in Nile tilapia breeding programs.

## Conclusions

The current study revealed similar short-range LD decay for three farmed Nile tilapia populations. The PCA suggested three distinct populations, whereas the admixture analysis confirmed that these three populations are highly admixed and are directly or indirectly related to the same GIFT strain origin. Based on the number of independent chromosome segments, at least 4.2 K SNPs might be required to implement GWAS and genomic prediction in the current Nile tilapia populations.

## Ethics approval and consent to participate

Nile tilapia sampling procedures is in process of approving by the Comité de Bioética Animal from the Facultad de Ciencias Veterinarias y Pecuarias, Universidad de Chile.

## Consent for publication

Not applicable

## Availability of data and material

For each each population, raw genotype data is available on the online digital repository Figshare, accession number 10.6084/m9.figshare.7581581.

## Authors’ contributions

GMY performed the analysis and wrote the initial version of the manuscript. AB contribute with discussion and writing. GC, MC and AJ performed DNA extraction. KC and JPL contributed with study design. JMY conceived and designed the study; contributed to the analysis, discussion and writing. All authors have reviewed and approved the manuscript.

## Acknowledgements

The authors are grateful to Aquacorporación Internacional and AquaAmerica for providing the Nile tilapia samples. We would like to thank to José Soto and Diego Salas from Aquacorporación International and Natalí Kunita and Gabriel Rizzato from AquaAmerica for their kind contribution with tilapia samples from Costa Rica and Brazil, respectively.

---

**Supplementary File 1.** Admixture clustering of the three Nile populations for K values ranging from 2 to 10. The animals are grouped by population and each individual is represented by a vertical bar. The gradient black lines delineate different populations under study.

**Supplementary File 2.** Linkage disequilibrium decay by physical distance estimated by chromosome linkage group (LG) for POP A.

**Supplementary File 3.** Linkage disequilibrium decay by physical distance estimated by chromosome linkage group (LG) for POP B.

**Supplementary File 4.** Linkage disequilibrium decay by physical distance estimated by chromosome linkage group (LG) for POP C.

