## Supplementary figures and images for "Genome-wide patterns of population structure and linkage disequilibrium in farmed Nile tilapia (*Oreochromis niloticus*)"

### Supplemental Figure 1

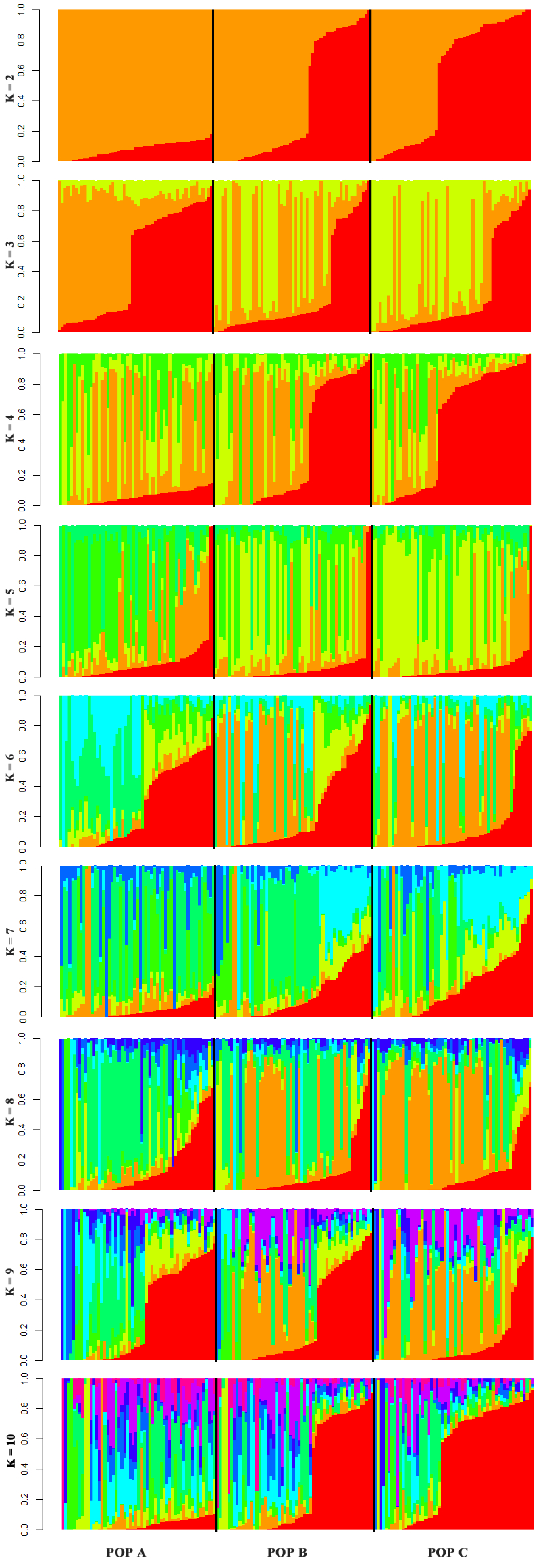

### Supplemental Figure 2

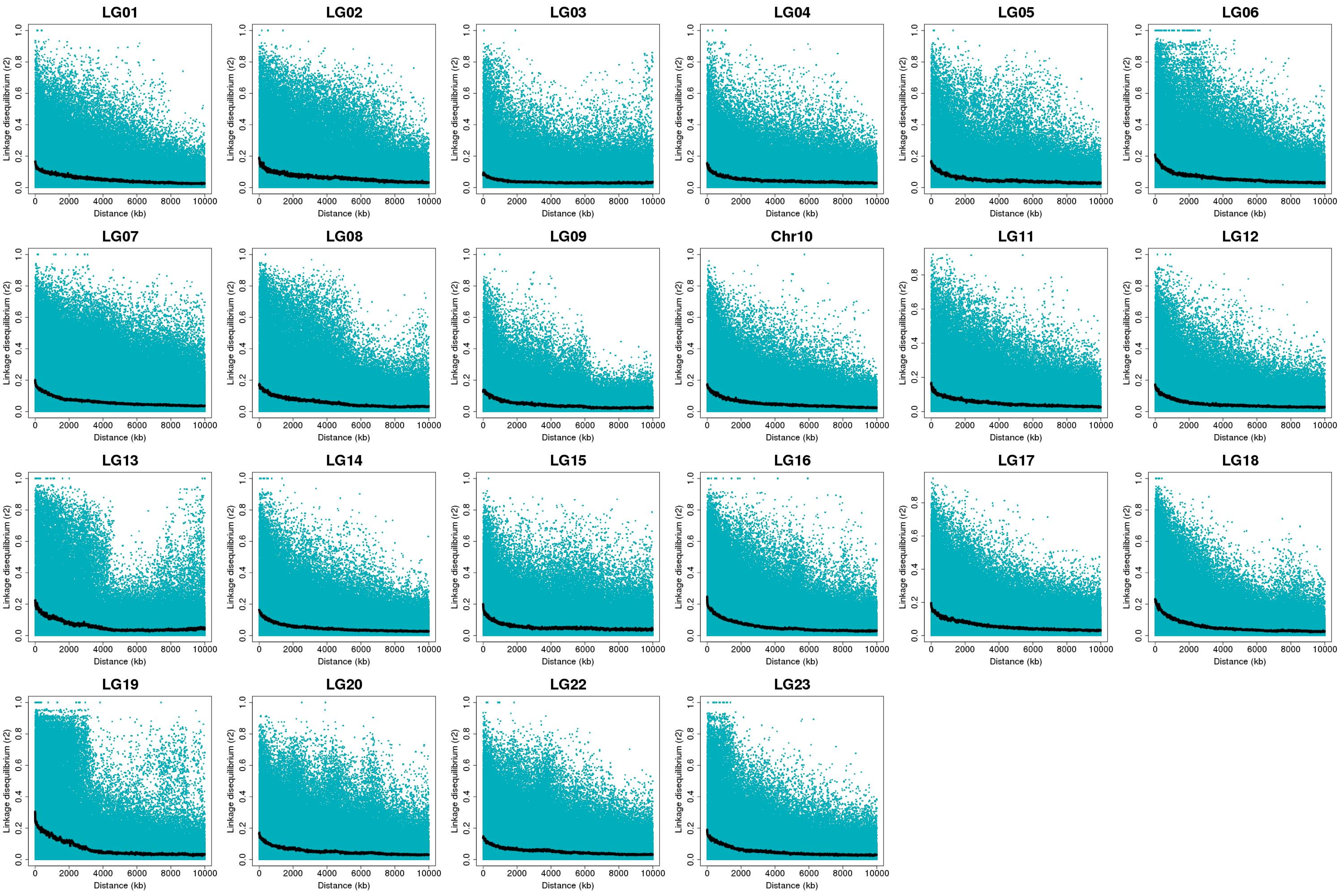

### Supplemental Figure 3

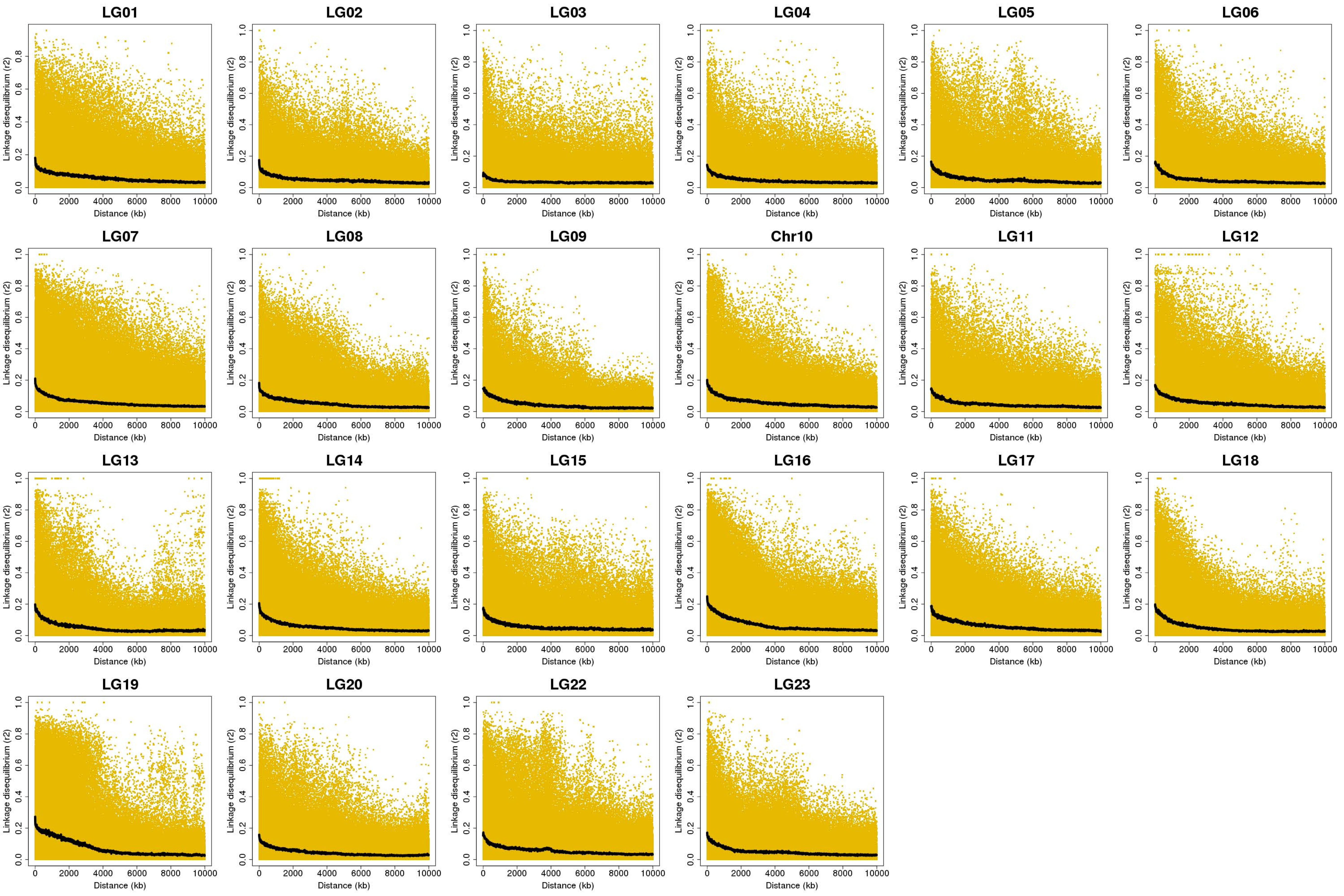

### Supplemental Figure 4

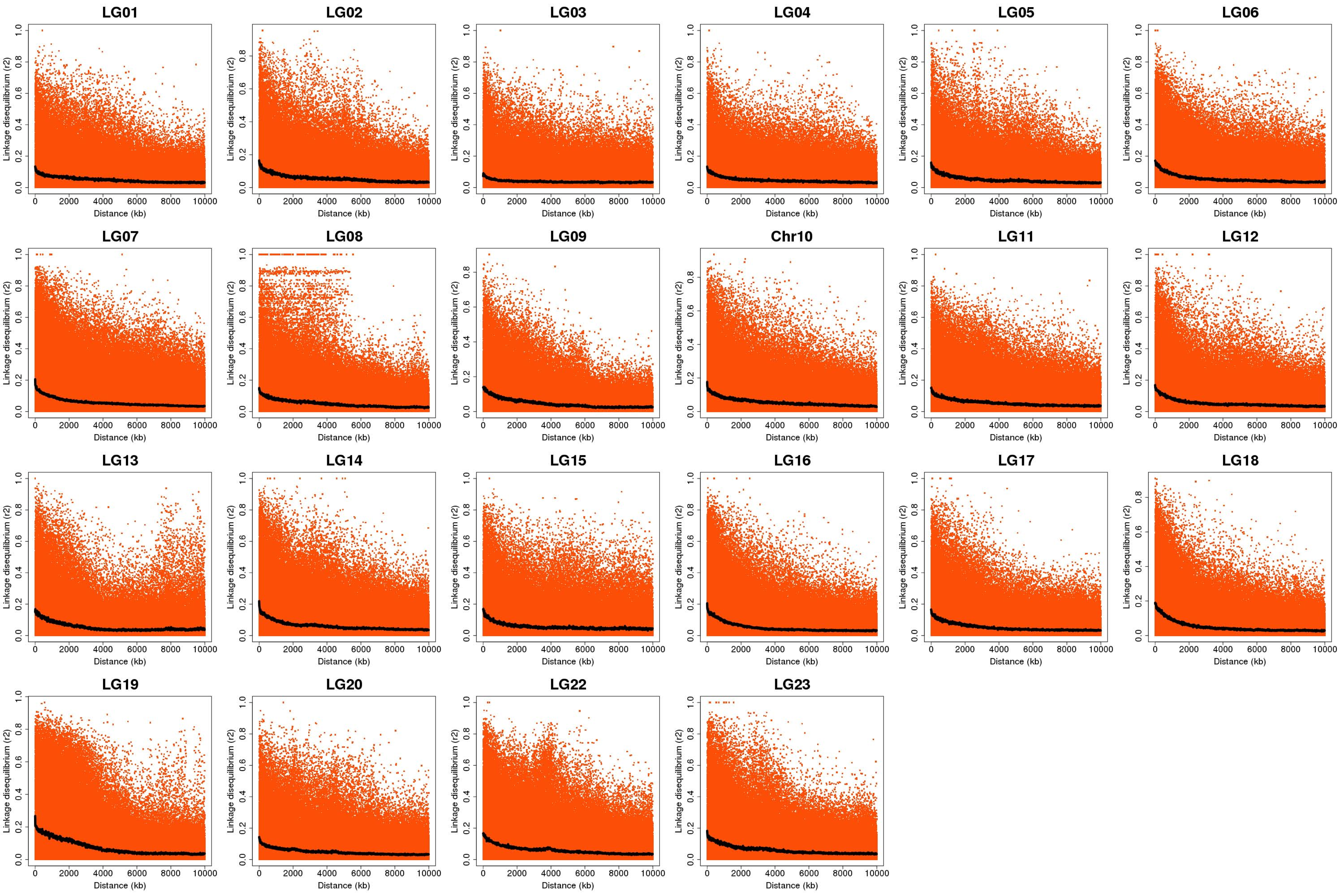
